# The dorsal thalamic lateral geniculate nucleus is required for visual control of head direction cell firing direction in rats

**DOI:** 10.1101/2024.05.06.592727

**Authors:** James S. Street, Kate J. Jeffery

## Abstract

Head direction (HD) neurons, signalling facing direction, generate a signal that is primarily anchored to the outside world by visual inputs. We investigated the route for visual landmark information into the HD system in rats. There are two candidates: an evolutionarily older, larger subcortical (retino-tectal) pathway specialised for coarse vision, and a smaller cortical (retino-geniculo-striate) pathway for higher acuity vision. We disrupted the cortical pathway by lesioning the dorsal lateral geniculate (dLGN) thalamic nuclei bilaterally, and recorded HD cells in postsubicular (PoS) cortex as rats foraged in a visual-cue-controlled enclosure. In dLGN-lesioned rats we found the expected number of PoS HD cells. Although directional tuning curves were broader across a trial, this was due to increased instability of otherwise normal-width tuning curves. Tuning curves were also poorly responsive to polarizing visual landmarks, and did not distinguish cues based on their visual pattern. Thus, the retino-geniculo-striate pathway is not critical for generation of an underlying, tightly-tuned directional signal, but does provide the main route for vision-based anchoring of the signal to the outside world, even when visual cues are high-contrast and low in detail.

**Key points:**

- Head direction (HD) cells indicate the facing direction of the head, using visual landmarks to distinguish directions
- In rats, we investigated whether this visual information is routed through the thalamus to visual cortex or arrives via the superior colliculus: a phylogenetically older and (in rodents) larger pathway, but not known for pattern vision.
- We lesioned the thalamic dorsal lateral geniculate nucleus (dLGN) in rats and recorded the responsiveness of cortical HD cells to visual cues.
- We found that cortical HD cells had normal tuning curves but these were slightly more unstable during a trial. Most notably, HD cells in dLGN-lesioned animals showed little ability to distinguish highly distinct cues and none to distinguish more similar cues.
- These results suggest that directional processing of visual landmarks in mammals requires the geniculo-cortical pathway, which raises questions about when and how visual directional landmark processing appeared during evolution.

## Introduction

The head direction (HD) system, most intensively studied in rodents, is a core part of the vertebrate navigation/memory system, without which a stable representation of the spatial environment could not be formed. It is also a potential, tractable model system for studying sensory integration and neural plasticity. HD neurons are distributed across several interconnected cortical and subcortical brain regions (Taube *et al*., 1990*a*; Taube, 2007). A single HD cell in one of these regions fires strongly when an animal faces a specific direction in the environment (referred to as its “firing direction”; FD), and as a population these cells encode all directions in the horizontal plane. The cells use information from familiar visual scenery in order to initiate their firing directions on entry into an environment (Taube *et al*., 1990*b*; Lozano *et al*., 2017), as well as to reorient the signal if it drifts. The present study investigated the route by which visual scene (landmark) information reaches the HD system.

There are, broadly speaking, two possible routes, corresponding to the two visual systems in the vertebrate brain (Figure 1). One is the evolutionarily older subcortical retino-tectal pathway, which connects the retina to the superior colliculus and is the largest visual pathway in rodents, carrying 90% of retinal projections (Ellis *et al*., 2016), and conveying low-detail information used for orienting movements. The other is the more recently evolved cortical retino-geniculo-striate pathway, which is small (only 25-50% of retinal output, (Linden & Perry, 1983; Martin, 1986; Ellis *et al*., 2016)), but has higher acuity and is used for processing fine details. Retinal inputs first reach the thalamic dorsal lateral geniculate nucleus (dLGN), and the target neurons then project to primary visual cortex (V1). Ever since the identification of the two systems in the 1960s it has been known that an intact retino-geniculo-striate system is not necessary for animals’ ability to respond to visual stimuli (see (Petry & Bickford, 2019) for review). Animals using only the retino-tectal pathway can avoid obstacles, learn visuospatial tasks, and recognise food (Humphrey & Weiskrantz, 1967; Dean, 1981; McDaniel *et al*., 1982), and humans with visual cortex lesions can do the same without being aware of their visual capability: a phenomenon known as blindsight (Cowey, 2010). Given that direction coding is an ancient and fundamental competence, possessed by animals such as insects and fish that diverged from mammals in evolution hundreds of millions of years ago (Knudsen, 2020), the question arises as to whether the direction system receives its visual inputs via the ancient retino-tectal or more modern retino-geniculo-striate pathway.

**Figure 1.**
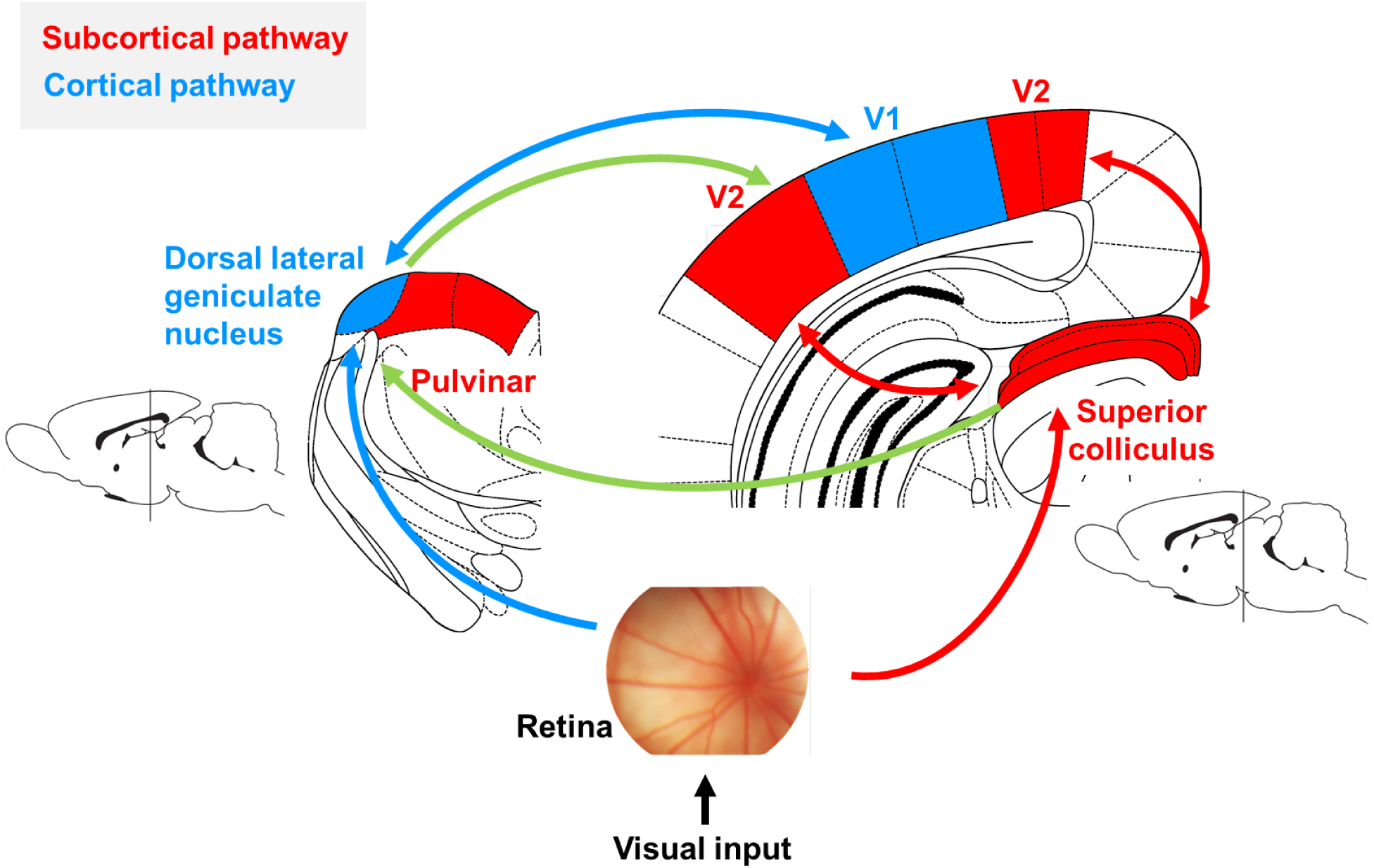
Two visual pathways.

The two pathways are not completely independent: for example, the lateral pulvinar of the thalamus (part of the retino-tectal pathway) enhances feature selectivity in mouse V1 by providing surround suppression (Fang *et al*., 2020), and the velocity selectivity of neurons in extrastriate cortices was impaired following collicular lesions (Tohmi *et al*., 2014). Nevertheless, the differential functions of these systems led us to ask the question of whether HD cells can use only the coarse vision of the retino-tectal pathway to mediate responsiveness to visual scenes, which would be consistent with growing evidence that the HD system is evolutionarily ancient. We did this by recording cortical HD neurons in rats in which the dLGN had been lesioned. Below, we outline in more detail the background to these questions.

### The head direction system and its visual influences

The head direction system comprises a network of brain regions, both cortical and subcortical (Figure 2). HD cells in these regions have in common that each cell fires intensely when the animal faces within a fairly restricted range of directions, typically around 90 degrees, and much less so, or not at all, in other directions. Different cells have different preferred firing directions (FDs) but by and large, manipulations(such as rotating a landmark) that cause one cell to reorient its firing direction affect all the other cells similarly, leading to a high degree of coherence across the population. These observations have given rise to the influential hypothesis that the cells intercommunicate in a so-called attractor network (Skaggs *et al*., 1995; Zhang, 1996; Angelaki & Laurens, 2020). By this model, the organisation of the network allows activity to be maintained within a restricted part of it at any particular moment, but also to smoothly move to a new set of cells when the animal turns its head. Because activity will eventually, if the animal keeps turning, return to its starting point (after 360 degrees of head turn), the network dynamic has an abstract ring topology (Chaudhuri *et al*., 2019) which is called a ring attractor. Evidence for a ring attractor has also been found in insects (Kim *et al*., 2017), where the anatomical organisation of the network is such that the cells form a physical ring (unlike in vertebrates where the physical location of the cells is random). The population of cells active at a particular moment is known in computational circles as a “bump”, and the property the network has of smoothly moving the bump from one part of the network to the next, instead of abruptly jumping, leads to its characterization as a continuous (as opposed to discrete) attractor.

**Figure 2:**
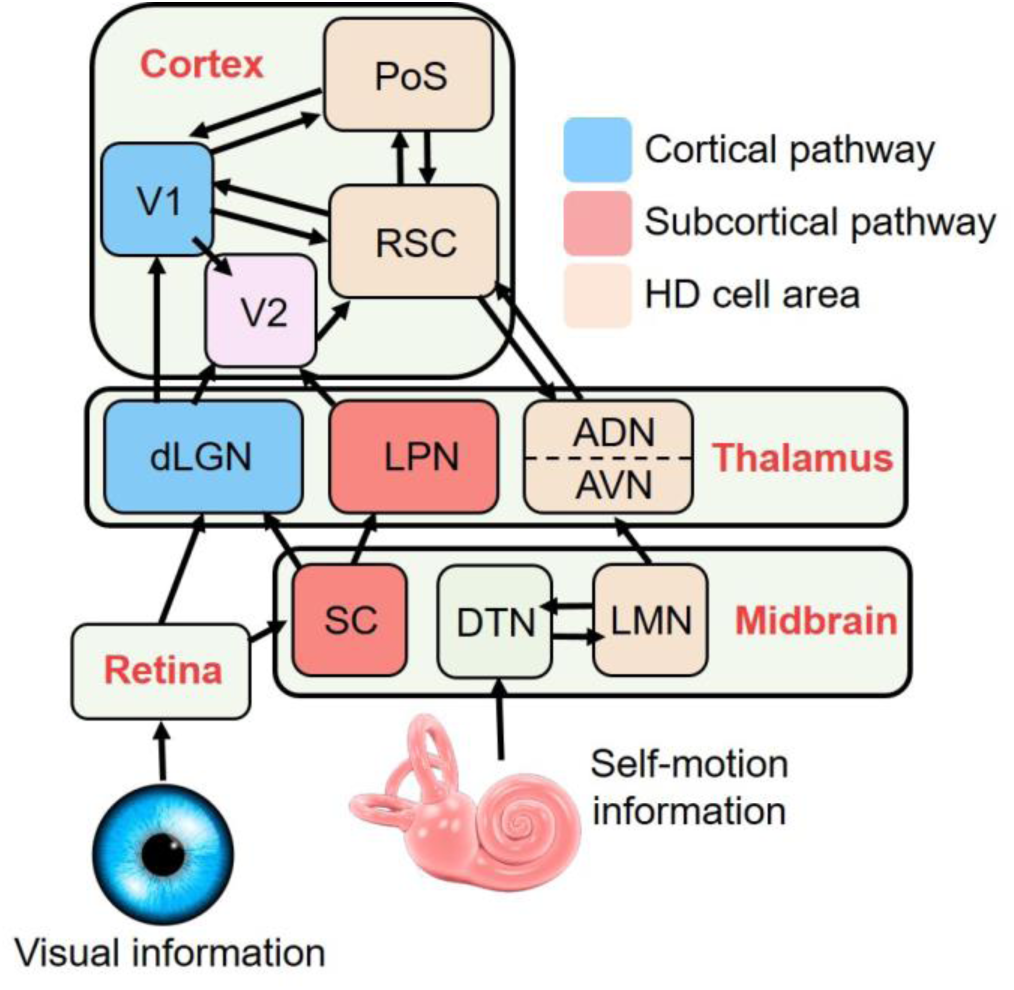
connectivity diagram of the interaction between visual and head direction systems.

HD cells are always active, but when an animal enters an environment the network needs some kind of initialization, so as to adopt a stable and reproducible orientation with respect to the environmental cues. In a familiar environment this orientation is retrieved from memory, reactivated by perception of the environment cues (Taube *et al*., 1990*b*). Even in rodents, this perception is mainly visual. If the environment is novel then the cells maintain the orientation they had at the moment of entry, and then learn the association of the new environment layout with this orientation (Taube & Burton, 1995). One way to demonstrate the influence of vision on this process is by recording HD neurons from animals exploring a cylindrical curtained-off environment, in which only visually contrasting cue cards break the visual rotational symmetry. In the absence of any such cues the symmetry is infinite: that is, the cells cannot use vision to establish a directional orientation. With a single visual cue the symmetry is completely broken and the cells, if they can detect the cue, can now unambiguously determine firing direction (Taube *et al*., 1990*b*; Sit & Goard, 2023; Ajabi *et al*., 2023). The same is true if there are two visibly different visual cues located on opposite sides of the arena (Lozano *et al*., 2017). However if the visual system cannot distinguish opposing cues then the environment’s rotational symmetry is twofold: each direction looks, to the system, exactly the same as its opposite. In this ambiguous situation, HD cells select one of the two possible orientations apparently at random, (Lozano *et al*., 2017), possibly determined by the network state at the moment of entry (Sharp *et al*., 1990). They then maintain that orientation for the remainder of the recording trial. Such maintenance is underpinned by the attractor dynamics discussed above.

The above two-cue paradigm provides a useful test of the visual acuity of the system, because HD cells can only adopt a consistent orientation relative to the cue-pair if they can distinguish the cues (Lozano *et al*., 2017). The present study used this paradigm to investigate the route by which visual environmental information reaches the cells. We made excitotoxic lesions of a key node in the geniculo-striate pathway, the dLGN, to deprive the system of high-acuity vision. We then recorded HD cells from the postsubiculum (PoS) to assess their firing characteristics. We chose PoS because it is the cortical HD region with the highest yield of HD cells, and is monosynaptically linked to V1. We asked the following questions: (i) Are there PoS HD cells in the expected numbers; (ii) Do they have the same basic characteristics as in intact animals; (iii) Do they rotate in response to rotations of the visual cues, and (iv) Do they distinguish the patterned visual cues? We find that disruptions of the geniculo-striate pathway, via bilateral excitotoxic lesions of the dLGN, severely impair visual landmark anchoring characteristics of HD cells. In lesioned animals, HD cells exhibit wider tuning curves and reduced directionality over the course of a trial, which can be explained as the average directional representation of an unanchored, drifting attractor. Accordingly, we found substantial impairment in the capacity of HD cells to rotate to follow rotations of prominent visual cues, with a greater impairment observed using cue cards requiring higher acuity.

## Methods

### Ethical approval

This study was approved by the UCL Animal Welfare Ethics Review Board (AWERB) and all licensed procedures were performed under the jurisdiction of a Project Licence granted to K.J.J. by the Home Office according to the Animals (Scientific Procedures) Act (1986), and European Communities Council Directive 86/609/EEC. All work was in line with the Journal of Physiology ethical principles and compliant with the journal’s animal ethics checklist.

### Animals

Eleven adult male Lister-Hooded rats (Charles River Laboratories Inc., UK; 300-600g) contributed data to this experiment. Animals were housed in a temperature and humidity controlled holding room on a 12h:12hr day:night cycle, containing 1h each of simulated dawn and dusk. Animals were singly housed following surgery. Until commencement of recordings, animals had access to food and water *ad libitum*; prior to commencing recordings animals were lightly food-restricted to target 90% of their free-feed weight.

### Surgical procedures: lesion and implantation

Lesions and electrode implantation took place in the same operation, using stereotaxic positioning. Isoflurane anaesthesia was used for induction (5%) and maintenance (1.5-3%) of anaesthesia, and depth of anaesthesia was monitored throughout by the pedal withdrawal reflex and respiratory rate. At the beginning of surgery, animals received subcutaneous analgesia (1 ml/kg carprofen 0.5% v/w).

The lesion process comprised bilateral excitotoxic lesions of the dorsal lateral geniculate nucleus (Figure 3A), made using injections of the excitotoxin *n*-methyl-D-aspartate (NMDA; 0.09M, Sigma-Aldrich) into the coordinates shown in Table 1. A glass pipette (0.66 mm diameter) filled with either NMDA or 0.9% saline was lowered through a craniotomy into the brain to each injection site, a small well was made (by repeatedly advancing the pipette tip +0.1 mm beyond the desired injection coordinate), and 0.25 μl of NMDA/saline was infused at a rate of 0.1 μl/min; these parameters are similar to those previously reported in the literature (Kirby *et al*., 2012). Four injections were made in the dLGN of each hemisphere, totalling 1 μl NMDA/saline injected per hemisphere (see Table 1). Where possible (when the dura was not damaged following craniotomies; 17/22 hemispheres injected), injections were made through intact meninges so as to minimise damage to the overlying cortex. 0.1 mm was added to the reported DV coordinate when injecting through the meninges. After each injection, the experimenter waited 5 minutes before retracting the pipette, to allow diffusion of the liquid into the brain and minimise reflux.

**Figure 3.**
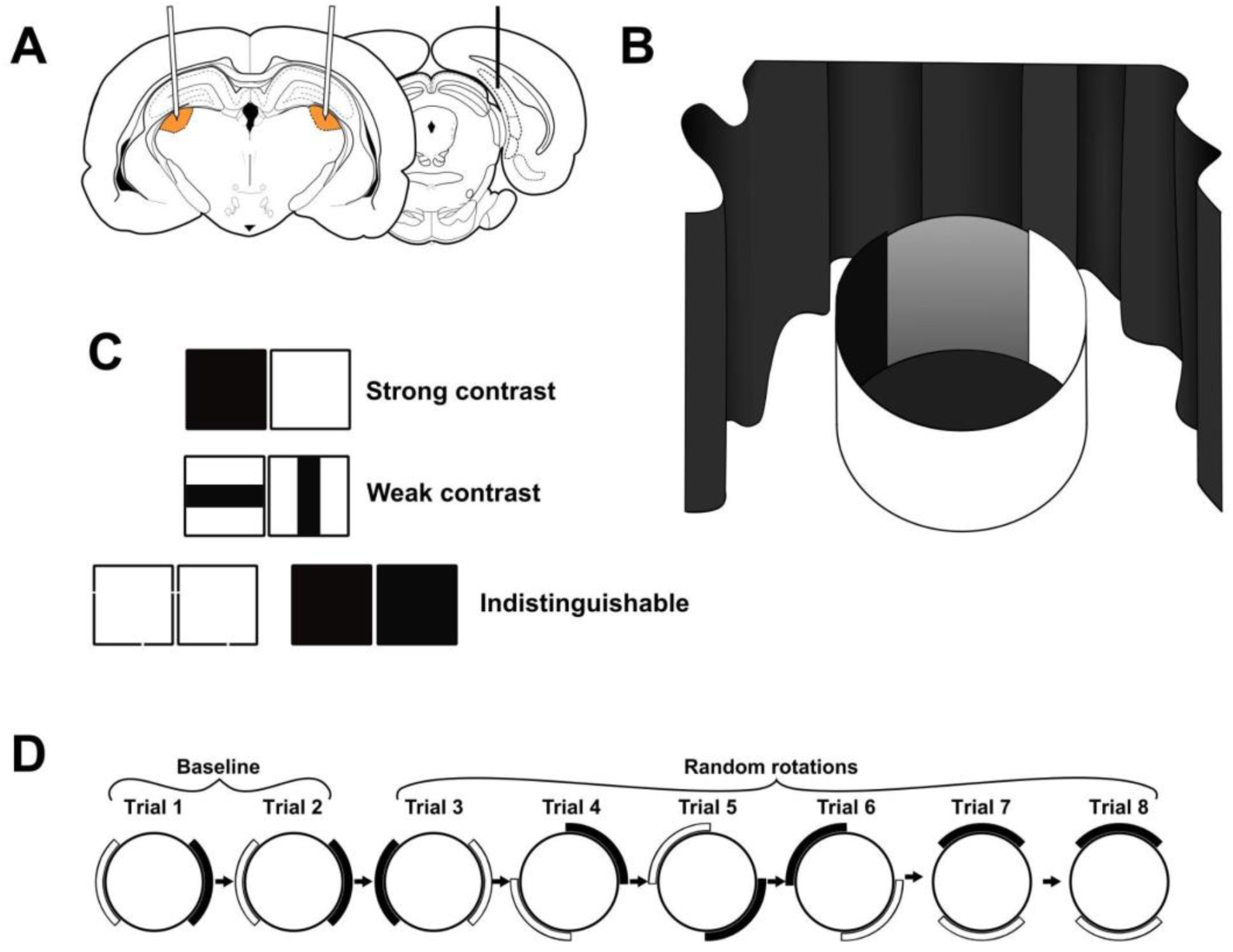
Summary of experimental design and recording protocol. A: Schematic of the surgical targets for dLGN NMDA injections (orange) and PoS tetrode implants (black line). B: Recording setup. The cylinder was 80 cm in diameter. C: Sets of cue cards used during recording sessions. D: Recording protocol for a single recording session. One session consisted of 8 trials, the first two of which were baseline trials in which the cylinder was not rotated, and the latter six in which the cylinder was rotated a random multiple of 45°.

**Table 1.** Coordinates used for bilateral dLGN injection and unilateral PoS implantation. AP and ML coordinates are given relative to bregma; DV coordinates are given relative to brain surface.

|  | Co-ordinates (mm) |  |  |
| --- | --- | --- | --- |
|  | AP | ML | DV |
| **dLGN injections** | - 3.7 | ± 3.2 | - 4.8 |
| | - 4.2 | ± 3.2 | - 5.2 |
| | - 4.8 | ± 3.4 | - 5.0 |
| | - 5.3 | ± 3.4 | - 5.0 |
| **PoS implant** | - 7.5 | ± 3.4 | - 2.0 |

After all injections were made, tetrodes implanted into the PoS (Figure 3A). These were attached to an Axona microdrive chassis and constructed from 90%:10% platinum-iridium alloy wire of 25 μm diameter (California Fine Wire). For the implantation procedure, a craniotomy was drilled above the PoS, the meninges removed, and the four tetrodes were lowered to the relevant coordinate (bregma −7.5 mm anteroposterior, +3.4 mm mediolateral, and 2 mm below brain surface) and cemented in place with dental cement.

At the end of surgery, animals received subcutaneous fluids (0.5-2ml Hartmann’s solution), and in some cases a single dose of diazepam (2.5 mg/kg intraperitoneal injection), in order to prophylactically reduce the risk of any seizure activity caused by the NMDA. Animals were given 3 days of postoperative analgesia (oral meloxicam), and allowed one week of recovery prior to recording.

### Electrophysiological recordings

For single-unit recordings, implanted drives were connected via a headstage and wire to a preamplification unit and digitiser unit (Axona Ltd, St Albans UK). Neural signals were amplified with a gain 5000-25000, and putative spikes (identified by amplitude crossings over a manually set threshold) were recorded and digitised at 48kHz, acquiring from −0.2 to +0.8 ms from threshold crossing. Animals were tracked using two infrared LED arrays mounted 8 cm apart on the headstage attached to the animal’s head. Position and head direction were extracted and aligned with the neural acquisition and saved for offline analysis.

To identify HD cells, initial screening recordings were performed in a box of dimensions 1 x 1 x 0.5 m or 1.2 x 0.6 x 0.6 m (length x width x height). The box had a single prominent cue card attached to the inner wall, and distal cues were provided by the objects around the room.

If an HD cell was identified, the animal was carried in an opaque box to a different room containing the experimental arena. The recording arena (Figure 3B) comprised an 80cm-diameter 50cm-tall grey-walled cylinder, and was surrounded by curtains to screen off visible distant cues. Two removable 50 x 50cm cue cards were attached to the inside wall of the cylinder, positioned at 180° apart, each subtending approximately 70° of arc. One pair of cue cards was used per session: either a black-white, a vertical-horizontal, a black-black, or white-white set of cards (Figure 3C). The two visually identical sets (black-black and white-white) were grouped together for analysis (referred to hereafter as the black-black cue condition). The first recording session for each animal used black-white cues, as this configuration was expected to exert the strongest cue control over the HD system (Lozano *et al*., 2017); in each subsequent session, the cues to be used were randomly selected.

Recording sessions (Figure 3D) consisted of 8 trials of 8 minutes each. Between trials 1 and 2 (‘baseline’ trials), the cylinder remained in its same orientation, but in trials 3 - 8 it was true-randomly rotated by *±*45°, *±*90°, *±*135°, or 180°. Before each trial, animals were lightly disorientated by the experimenter by rotation, prior to being placed in the cylinder. Sessions were analysed if at least 4 trials were successfully recorded.

Animal position and head direction during exploration of the environment was monitored by tracking two infra-red LEDs mounted onto the headstage on the animal’s head. For each trial, a number of behavioural metrics were extracted from the reconstructed path of the animal: total distance travelled, average linear speed, total angular travel distance (sum of absolute angles), and average angular speed. We also calculated the total angular travel displacement (sum of signed angles) for each animal in each trial, to assess for the presence of turning bias.

Thigmotaxic (wall-hugging) behaviour was assessed using the animal’s tracked path. The smallest ellipse containing all tracked points was fitted to provide an estimate of the location of the cylinder walls. A smaller ellipse of the same eccentricity was then constructed around the centre of the larger ellipse containing half of its area; this smaller ellipse provided a geometrically defined ‘centre-half’ of the cylinder. The fraction of time spent in the smaller ellipse provides an inverse measure of thigmotaxis.

### Spike clustering and cell isolation

Recorded electrophysiological data were analysed off-line. Presumptive neuronal spikes were clustered using the clustering software Tint (Axona Ltd., St Albans, UK) and KlustaKwik v3.0 (Harris *et al*., 2000), followed by manual refinement of clusters in the graphical interface. Cells were excluded if their cluster contained substantial noise across the session (if >1% of spikes occurred within a 2ms refractory period in the autocorrelogram). L-ratio and isolation distance (Schmitzer-Torbert *et al*., 2005) were derived for each HD cell cluster to provide cluster quality metrics. The refined clusters were saved and imported into MATLAB (2015a or 2019b, Mathworks Ltd., Natick, MA) for further analysis using the CircStat package (Berens, 2009).

### HD cell classification

Head direction samples were binned into 60 bins of 6° width, and the trial-averaged firing rate in each bin was calculated as the number of spikes in each bin divided by the total directional dwell time. The resulting histogram was smoothed using a boxcar kernel of 5 bins width to yield the head direction tuning curve for that cell, on which subsequent analyses were performed.

Putative neurons were classified as HD cells if they fulfilled all three of the following criteria in trials 1 and 2 of the recording session: (i) exhibited a peak firing rate of >1Hz, (ii) a Rayleigh vector length of the tuning curve of *R* > 0.3, and (iii) *R* > 99th percentile of a control distribution derived from a shuffle procedure. This shuffle was calculated by repeatedly circularly time-shifting the spike-train by a random time between 20s and 460s, and then recomputing the tuning curve and Rayleigh vector for each time-shifted spike-train. This was done 10000 times for each cell.

### Cell firing characteristics

For each recorded HD cell, we calculated the neuron’s (i) peak firing rate (defined as the value of the bin with greatest firing rate in the smoothed tuning curve), (ii) preferred firing direction (the direction of the peak firing rate bin in the smoothed tuning curve), (iii) tuning curve Rayleigh vector length, (iv) tuning width (the circular standard deviation of the tuning curve), (v) and directional information content (Skaggs *et al*., 1992).

### Drift analysis

To evaluate the drift of HD cell tuning curves over a trial, we initially divided the trials into two halves of 4 minutes each, and constructed a tuning curve for each half. The difference in firing directions of the tuning curves from the first and second trial halves provided an initial estimate of within-trial HD cell stability.

To further characterise any potential drift, we looked for evidence of instantaneous widening of HD tuning curves. To do so, we identified all events during a recording trial where the animal made a single head-turn that spanned the entire trial-averaged tuning curve of the HD cell (defined as ±90° either side of the cell’s peak firing direction). For each of these events, we calculated the angular difference between the first and last spike fired by the HD cell during the trajectory. If the HD cell was maintaining a precise tuning curve that drifted over time, the mean angle difference would be smaller than the trial-averaged tuning curve width. Conversely, a non-drifting cell would have an angle difference on individual head-turns that is similar to the whole trial-averaged width. This analysis provides a method to identify drifting HD cells without presupposing the rate or direction of the drifting behaviour.

### Attractor coherence

To investigate whether the HD attractor remained coherent through recordings, we assessed the intra-trial stability of co-recorded HD cells. This analysis was complicated by the drift in the signal, seen in lesioned animals, which could produce a spurious loosening of coupling if comparisons were made over too long a timescale (for example, the first and second halves of a trial). To overcome this, we looked over much shorter timescales using a boxcar procedure that compared firing directions in minute-long epochs, with each epoch shifted 12s along from the previous one. Thus, in trials where >1 HD cell was recorded, a 60-second time window was step-wise advanced across the recording in 12-second increments, and the mean spiking direction calculated for each cell in that window and then tuning curves pairwise compared with a Rayleigh vector of the angular differences.

### Landmark anchoring analysis

This analysis computed the firing direction (FD) shift between trials that would be expected if the cell were following the visual cue, and then determined the actual deviation of the FD from this shift. First, the rotation angle of the cylinder between trials was subtracted from each cell’s FD in each trial, so as to express each FD relative to the local visual cues. Then, for each session we selected a trial to use as a baseline against which to reference the FD shifts for all the HD cells in that session (see Figure 4). As cells may sometimes anomalously shift their relative FD in a single trial but revert back in later trials, we selected the trial containing the FD around closest to the session average. Specifically, we selected the trial that maximised the following metric, where *θ_j,i_* is the FD of cell *j* in trial *i*:

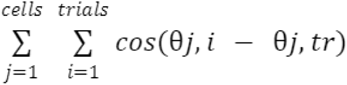

**Figure 4:**
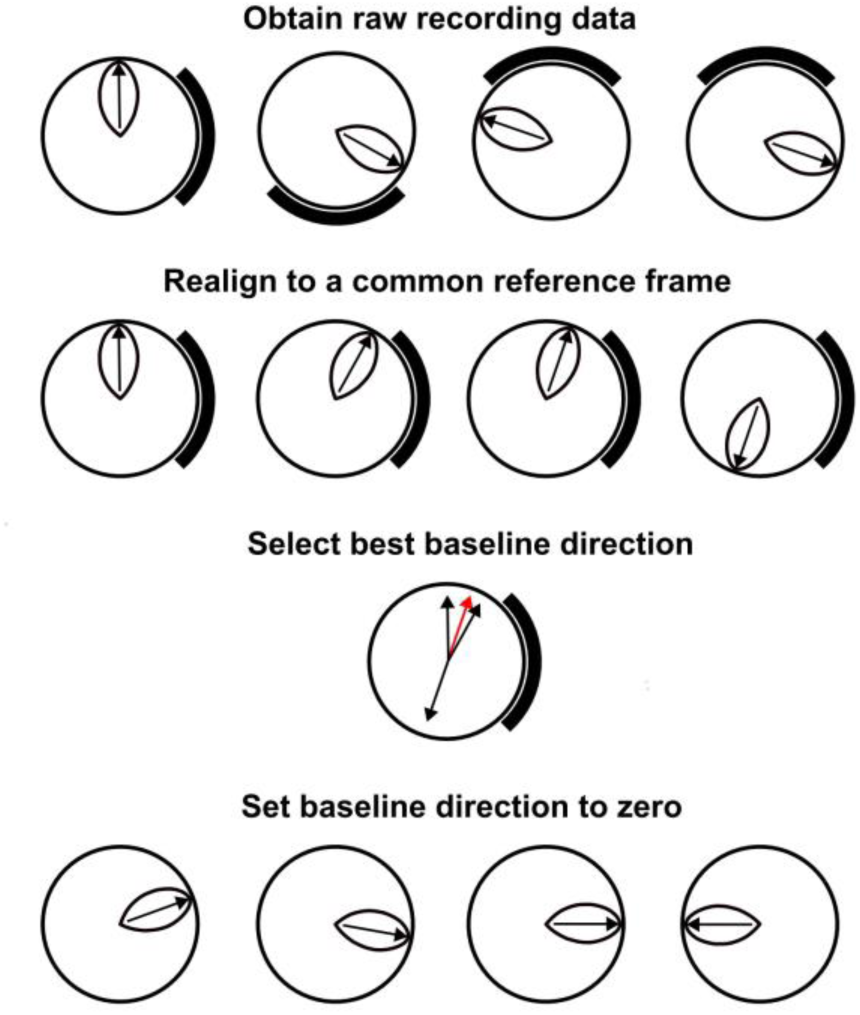
Landmark anchoring normalisation method. Schematic illustrating the methodology used to select a baseline trial against which firing direction shifts were normalised. In this set of four simulated trials, the HD cell maintains a consistent representation of direction in three of four trials; in trial 4, the cell shifts by 180°. Initially, all recorded tuning curves were realigned to the local reference frame of the cylinder. A baseline trial was then selected (red arrow) that best represented the most consistent directional representation maintained by the co-recorded cells during the recording session. All trials were subsequently normalised to this baseline, which was removed from the dataset prior to further analysis.

FDs of all cells in the baseline trial were subtracted from their cell FDs in all other trials, generating an estimate of FD shifts used for further statistical analysis. The selected baseline trial was removed from further analysis as its FD deviation was by definition zero.

Note that the same baseline trial was selected for all HD cells co-recorded during the session. If HD cells subtended a consistent angle relative to the cue cards, this would normalise the majority of FD shifts across all cells to near 0°. Any deviation from the expected angle relative to the cues would be reflected by greater FD shifts after normalisation.

As HD cells are believed to rotate coherently (Yoganarasimha & Knierim, 2005; Yoganarasimha *et al*., 2006), in each trial we averaged the FD shift across all cells recorded in that trial, providing a per-trial population estimate of heading direction which could be combined across sessions and animals.

### Animal-averaged anchoring metric

As different animals contributed different numbers of sessions, we derived a single summary metric for each animal in each cue condition to represent landmark anchoring precision in that animal. For this, we used a summed cosine transform of all FD shifts, normalised to the number of shifts recorded:

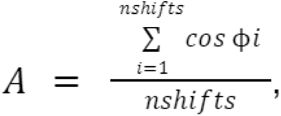

Values of A close to 1 indicate good average anchoring to that cue configuration, whereas values closer to 0 or below indicate poor anchoring.

### Statistical analysis

Assessment for significant non-uniformity of FD shifts was performed using the following circular statistics (Berens, 2009)): the Rayleigh test (which assesses unimodal deviations from uniformity in a circular distribution), the V test (which assesses deviations in a circular distribution towards a predicted direction), and the Kuiper test (which assesses whether the circular cumulative distributions of two groups are significantly different. Linear distributions are plotted as raincloud plots (Allen *et al*., 2019) using custom-written MATLAB functions. Linear data were tested for normality using the one-sample Kolmogorov-Smirnov test, with statistical comparisons performed using parametric or non-parametric methods as appropriate.

All statistical tests are two-tailed (alpha = 0.05) unless stated otherwise; all figures are reported as mean ± standard deviation (for parametric data) and median ± interquartile range (for nonparametric data) unless stated otherwise. In all figures * = significant at the 0.05 level, ** = significant at the 0.01 level, *** = significant at the 0.001 level.

### Histology

At the end of the experiment, rats were anaesthetised under isoflurane (5%) and killed with an intraperitoneal overdose of pentobarbitone, before being perfused transcardially with 0.9% saline followed by 10% neutral-buffered formalin. The brain was extracted and kept in a formalin solution for a minimum of 24 hours, and subsequently transferred to 30% sucrose by weight in phosphate-buffered saline for dehydration. Brains were frozen upright (Tissue-Tek OCT, Sakura FineTek) using a cryostat (CM1850 UV, Leica Biosystems, UK) to approximately −21°C, sectioned into 30-50 µm coronal slices, and stained using Cresyl violet for assessment of lesion extent and electrode placement. Light microscopy images were scanned using a Leica inverted epifluorescence microscope (DMi8, Leica Biosystems, UK) and acquisition software (LAS X, Leica Biosystems, UK).

Lesion extent was quantified by first selecting a representative brain slice for each slice in the rat brain atlas between −3.36 mm and −5.40 mm, that was considered to best fit the atlas plate (Paxinos & Watson, 2006). This was assessed by the experimenter through reconstruction of the known distance between adjacent brain slices, and by references to prominent landmarks. A polygonal outline was then traced around the lesion and around the dLGN using the open-source histological image analysis software ImageJ (Schindelin *et al*., 2012). The lesion extent was estimated by tracing the region within thalamus containing no visible neuron nuclei, while the dLGN was delineated using either changes in cellular architecture where visible (Evangelio *et al*., 2018) or (as was frequently the case in Lesion animals) from the relative positioning of local landmarks such as the thalamic border and third ventricle.

## Results

We recorded PoS HD cells from both groups of animals: 51 sessions of 406 trials in 6 Lesion animals, and 36 sessions of 282 trials in 5 Sham animals. Recordings were made while the animals freely foraged in a cylinder with two prominent cue cards attached 180° apart on the walls). The use of two cues enabled us to assess whether HD cells could not just detect the presence of a cue card but could also discriminate the visual content of the cards. Visually identical cue cards in some trials enabled us to decouple anchoring to tactile or olfactory components of the cue cards from anchoring to their visual content.

In what follows, we report results at the trial level (taking all trials from all animals) and at the animal level (averaging across trials within each animal).

### Histology

#### Lesion extent

Lesion extent (Figure 5) was estimated by analysis of Nissl-stained coronal slices taken through the extent of the dLGN. In all lesioned animals, a substantial (> 50%) portion of the dLGN was destroyed following NMDA injection. As expected, the proportion of damaged dLGN was greater in Lesion animals than Sham animals (Figure 5B and C) for representative slices through animals), although there was variation in lesion extent within the lesioned group. Figure 5D shows the estimated percentage of damaged dLGN across the 20 slices in each animal, assessed along the AP axis, for the left and right hemispheres. In general, lesions extended well across the AP axis of the dLGN, but in some animals the lesion did not extend to the far anterior/posterior poles of the nucleus. The total estimated %-area of lesioned dLGN for each animal is summarised in Table 2 alongside summary electrophysiological metrics (see Results - Electrophysiology).

**Figure 5:**
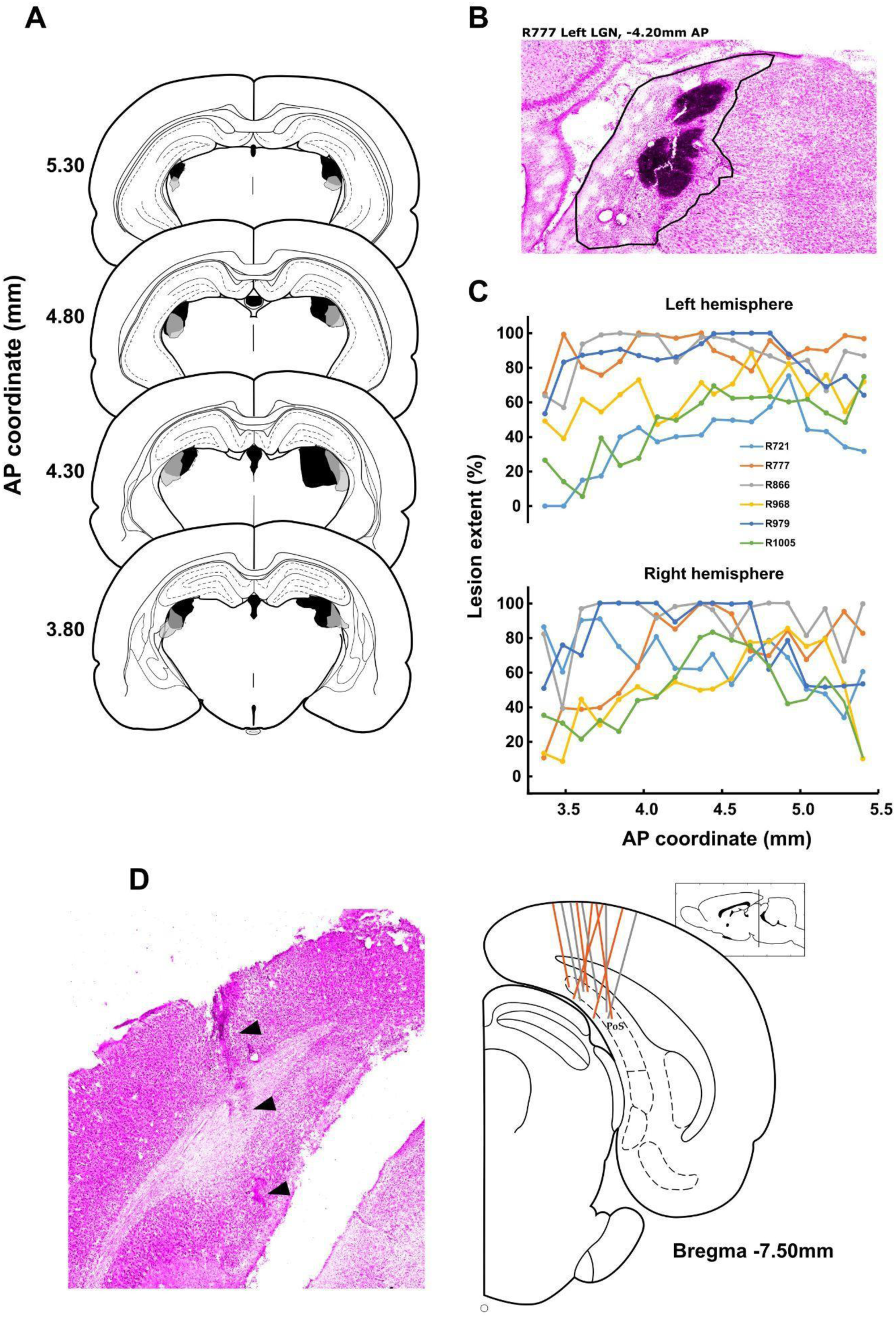
Histology. A-C Assessment of lesion extent. A: Schematic of the largest (grey) and smallest (black) lesions by % dLGN damaged. B: Example brain slice depicting left dLGN, with the lesion extent traced (black line). C: Lesion extents as a function of AP location for each individual animal. D: Left - Example brain slice depicting implant site in postsubiculum with electrode track (black arrows). Right - composite diagram of all electrode tracks, projected onto the same atlas slide. Inset shows the AP location of the section.

**Table 2.**
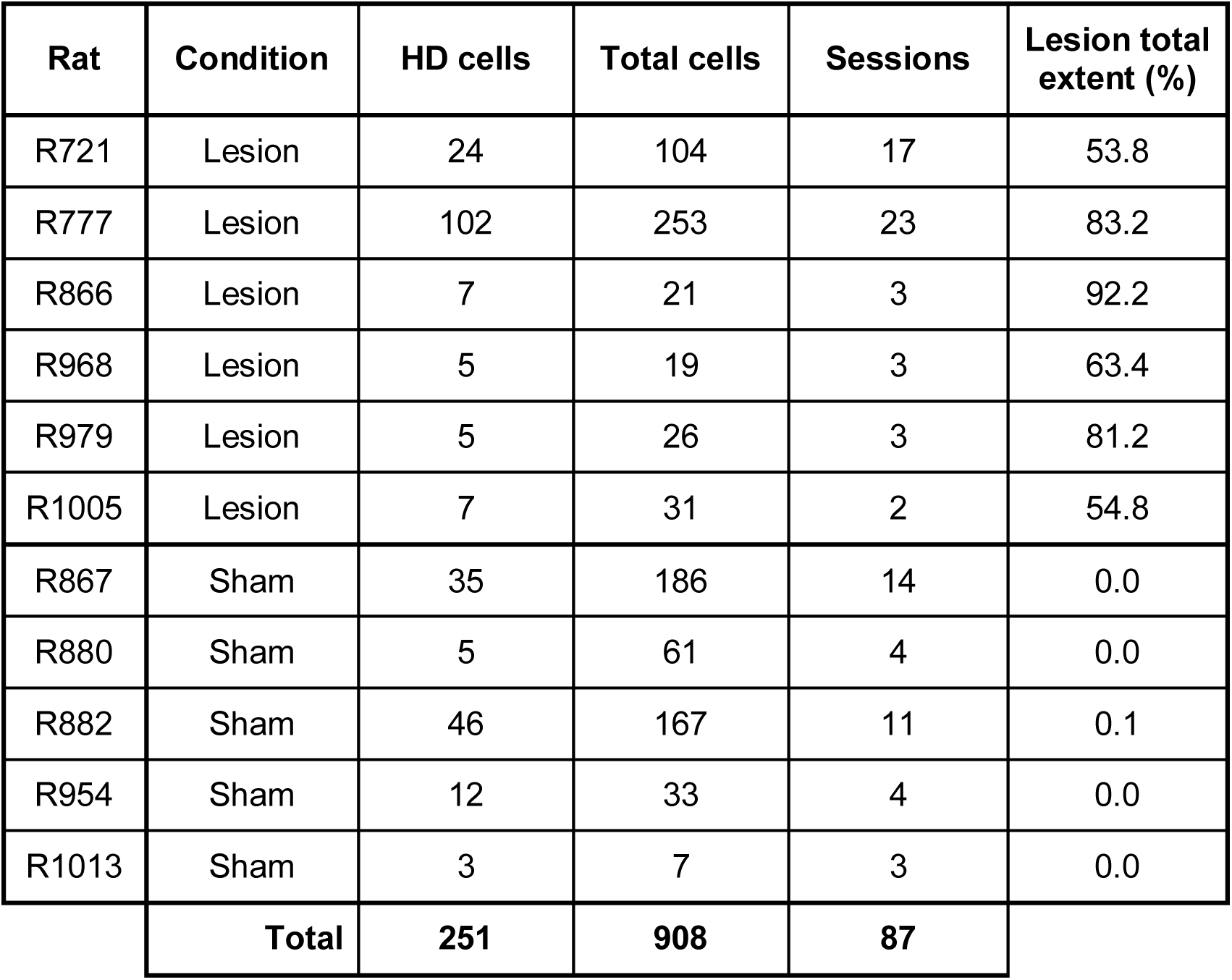
Numbers of cells, HD cells and sessions recorded from each animal.

#### Electrode tracks

The location of the recording site was determined by examination of the location of the electrode tracks (Figure 5D) and their apparent terminations, validated by estimation of the depth taken from a count of the microdrive screw turns since the implant depth. In all animals, evidence was seen of electrode tracks targeted successfully towards PoS.

### Behaviour

#### dLGN lesions mildly increased exploratory behaviour

As the spatial tuning of HD cells can be modulated by movement, we first compared the exploratory behaviour of Lesion animals relative to controls in the open field environment. We observed a significant difference in both linear and angular distance travelled over all trials between Lesion animals and Sham (median Lesion distance per trial: 25.1 ± 10.6 m, n=406 trials; median Sham session: 18.4 ± 7.9 m, n = 282 trials; Wilcoxon ranksum: z = 10.7, p < 10^-25^; Figure 6A), although this did not remain significant when averaged across all trials in each animal (linear: Lesion: 22.1 ± 4.6 m, n = 6 animals; Sham: 18.3 ± 6.7 m, n = 5 animals; Wilcoxon ranksum, p = 0.43; angular: Lesion: 2.62 ± 0.58 x 10^6^ °; Sham: 2.28 ± 0.85 x 10^6^ °; Wilcoxon ranksum, p = 0.93) travelled per trial. Thus, although there may be a mild increase in exploratory activity following dLGN lesions, lesions did not result in patent hyperactivity in animals.

**Figure 6:**
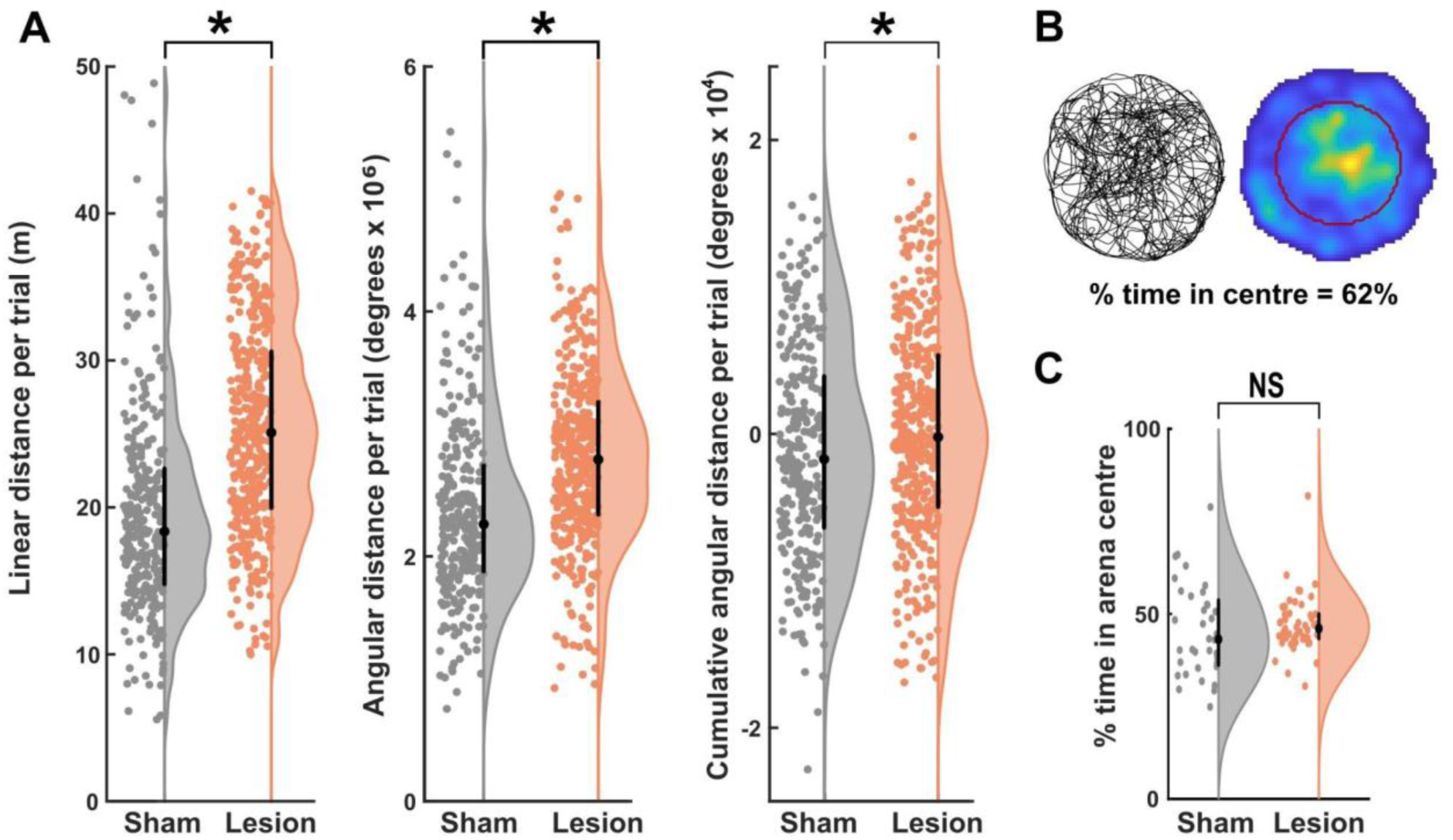
Behavioural characteristics of Sham and Lesion animals. A: Average movement statistics for individual animals. Left: Mean linear distance travelled by each animal per trial. Centre: Mean (absolute) angular distance travelled by each animal per trial. Right: Mean (directional) angular displacement travelled by each animal per trial. B: Thigmotaxis analysis example. Left: path of a Lesion animal during one trial. Right: Heat map of animal occupancy in the two-cue cylinder during the same trial, with the centre half of the cylinder highlighted (red line). C: Average time spent in centre half of the two-cue cylinder across all sessions.

We assessed whether Lesion animals might exhibit a turning bias during exploration by calculating the absolute difference in angle turned through clockwise (positive) versus counterclockwise (negative) in each trial.

We saw some evidence of a minor difference in turning bias at the individual trial level (median proportional turn bias, Lesion: 0.19 ± 0.26%, Sham: 0.23 ± 0.30%, Wilcoxon ranksum, z = −2.18, p = 0.0294), however these values were close to 0% in both Lesion and Sham animals, and this significance did not persist at the animal level (median Lesion animal turn bias: 0.19 ± 0.092%, median Sham animal turn bias: 0.19 ± 0.096%; Wilcoxon ranksum, p = 0.762).

We also considered whether visual deficits may lead Lesion animals to exhibit more thigmotaxis, remaining near the walls of the cylinder where they could be more sure of their location. To analyse this we divided the two-cue cylinder into inner and outer zones of equal area, and determined the proportion of time spent in each (Figure 6B. There was no difference in occupancy of the central zone between Lesion and Sham animals either at the session level (Figure 6C; median Lesion proportion in centre: 0.46 ± 0.07, median Sham proportion in centre: 0. 43 ± 0.17; Wilcoxon ranksum: z = 1.22, p = 0.22) or animal level (median Lesion animal proportion in centre: 0.46 ± 0.12, median Sham animal proportion in centre: 0.48 ± 0.15; Wilcoxon ranksum, p > 0.9), with values consistently close to 0.5, indicating that both groups of animals explored the cylinder uniformly and with no evident thigmotaxis. Taken together, these results indicate that Lesion and Sham animals exhibited broadly similar open field behaviour.

### Electrophysiology

#### Basic HD cell characteristics - cluster quality and firing rates did not differ

We recorded 251 HD cells from a total of 908 putative units in 11 rats (Table 2). Of these, 150 HD cells were recorded from 6 Lesion animals, and 101 HD cells were recorded from 5 Sham animals. There was no difference in the number of HD cells encountered in the two groups (average of 25.0 and 20.2 in Lesion and Sham groups respectively, NS). We saw no difference in HD cell cluster quality between Lesion and Sham groups (median L-ratio: Lesion: 0.23 ± 0.64, Sham: 0.32 ± 0.71; Wilcoxon ranksum, z = −0.87, p = 0.38, n = 251 clusters). Unexpectedly, clusters were significantly more likely to fulfil HD criteria in Lesion animals (Lesion: 33.0%, Sham: 22.3%; χ^2^ test of proportions with Yates correction = 12.69; p < 0.0005). Median HD cell peak firing rates and directional information content (Figure 7A) in Lesion and Sham animals did not differ (median peak firing rate, Lesion: 3.17 ± 4.20 Hz, Sham: 3.42 ± 5.37 Hz; Wilcoxon ranksum, z = −0.75, p = 0.45; median directional information, Lesion: 0.61 ± 0.88 bits/spike; Sham: 0.69 ± 1.25 bits/spike; Wilcoxon ranksum, z = −1.44, p = 0.15).

**Figure 7:**
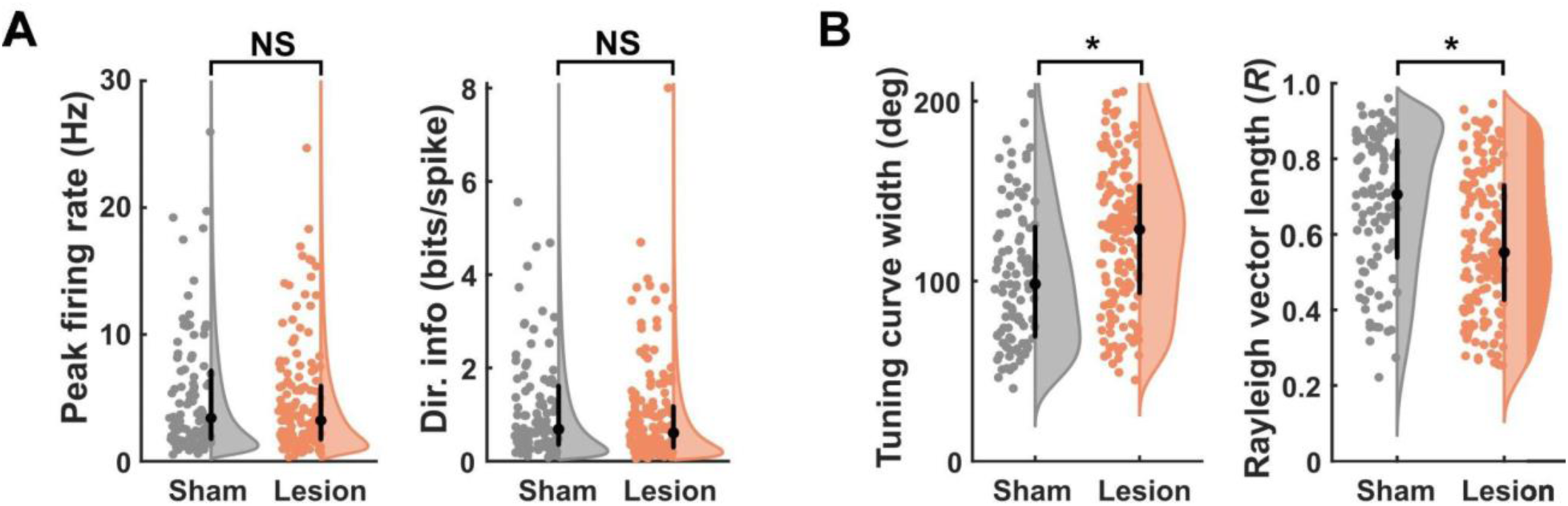
HD cell firing characteristics in Sham and Lesion animals. A: Firing rate and directional information content did not differ between Sham and Lesion animals. B: Tuning curves in Lesion animals were broader, as shown by the increased tuning curve width and slightly decreased Rayleigh vector length.

#### Broader HD cell tuning curves in dLGN-lesioned animals

HD cells in Lesion animals displayed broader firing (Figure 7B) as shown by wider mean tuning curves (mean tuning width, Lesion: 124 ± 40°, Sham: 103 ± 38°; t(249) = 4.24, p < 10^-4^) and smaller Rayleigh vector lengths (mean R vector, Lesion: R = 0.58 ± 0.19, n = 150, Sham: R = 0.68 ± 0.19, n = 101; t(249) = −4.13, p < 10^-4^; Figure. 7B). These results remained at significance at the single animal level (mean tuning width: t(9) = 2.51, p = 0.033; mean R vector: t(9) = −2.56, p = 0.031). Together these results indicate that HD cells maintained a less precise representation of direction following dLGN lesions.

#### HD cells in dLGN-lesioned animals drifted more across a trial, but coherently

We investigated whether the broader tuning curves in Lesion animals might be due to greater drifting, across a trial, of normal-width tuning curves (for example trials, see Figure 8A). Drift is plausible because previous evidence suggests HD cells exhibit drift of their tuning curves when animals are entirely deprived of vision (Goodridge *et al*., 1998; Butler *et al*., 2017; Asumbisa *et al*., 2022; Ajabi *et al*., 2023). To do this, we first took a broad-brush approach and examined how much HD cells changed their firing directions between the first and second halves of each trial. Lesion HD cells displayed greater absolute half-trial FD shifts averaged across sessions (median FD half-trial shift, Lesion: 29.3 ± 38.3°, Sham: 7.5 ± 14.6°; Wilcoxon ranksum, z = 7.54, p < 10^-13^). Consistent with this, the drift rate of HD cells, estimated across the entire trial using circular-linear regression, demonstrated a significantly greater absolute drift rate in Lesion HD cells than Sham over all trials (median drift rate, Lesion: 7.00 ± 14.0 °/min, Sham: 4.33 ± 6.67°/min; Wilcoxon ranksum, z = 8.66, p < 10^-17^, Figure 8B), and when averaged over animals (Lesion: 7.78 ± 1.60 °/min, Sham: 4.57 ± 1.39 °/min; t(9) = 3.51, p = 0.007).

**Figure 8:**
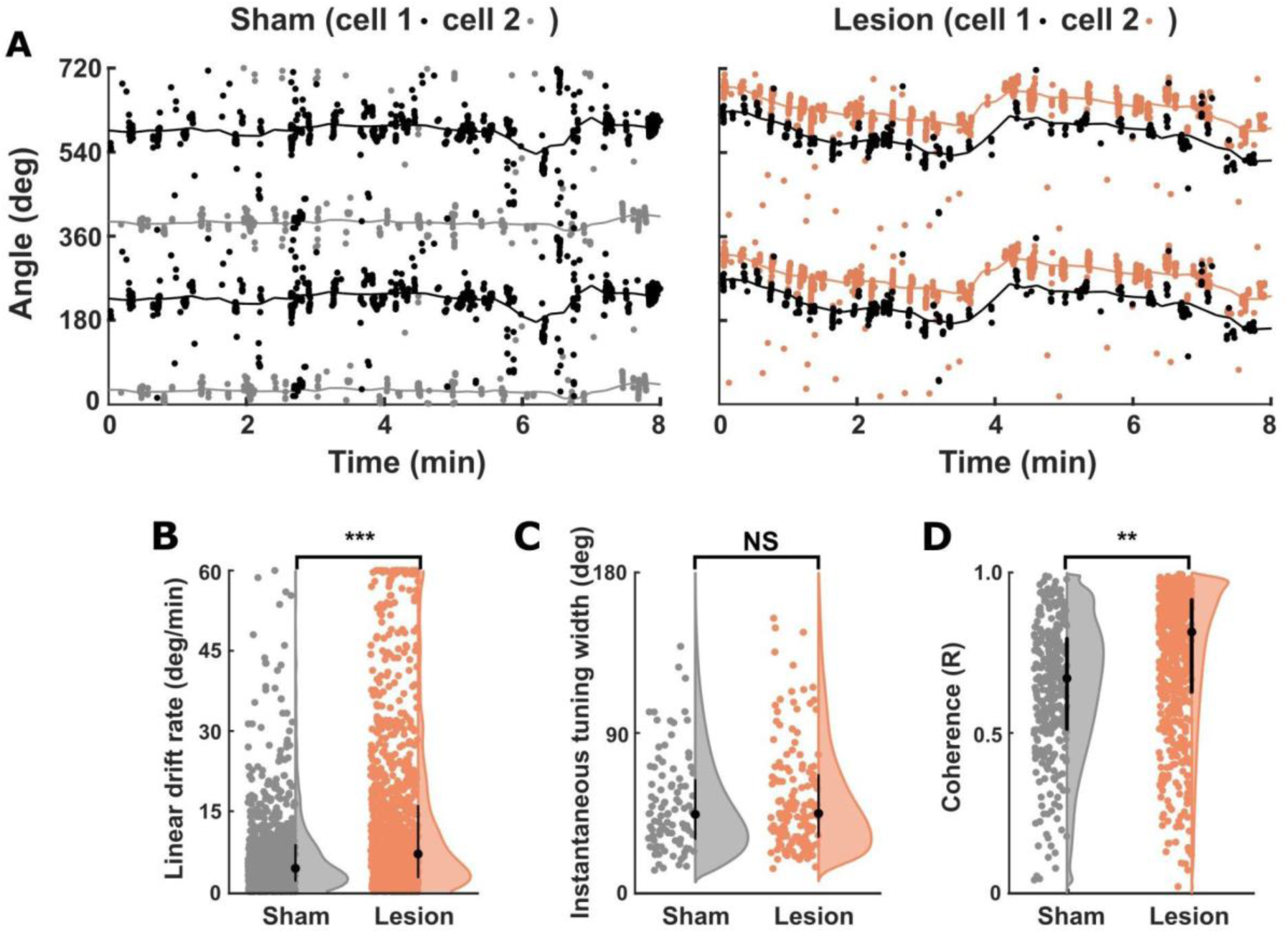
HD cell drift. A: Raster plot examples for two trials, each with two co-recorded HD cells, displaying the time and head direction at each spike. Left: Sham trial with two HD cells demonstrating high coherence and little evidence of drift. Right: a Lesion trial with two HD cells demonstrating high coherence but also significant drift throughout the trial. B: Distribution of fitted linear drift circular rate of HD cells in each trial, for Sham and Lesion HD cells. Lesion HD cells showed significantly increased median drift rates relative to Sham animals. C: No difference was seen in the instantaneous tuning widths of Sham and Lesion HD cells. D: Lesion HD cells remained significantly more coherent within individual trials than Sham HD cells. B-D significance levels shown for the per-cell analyses.

These analyses underestimate the true drift rate, since drift can occur in both directions and may cancel out across a whole trial. We thus undertook a more detailed analysis to estimate the tuning curve width over much shorter timescales. To do this we isolated epochs from the spike trace when the animal had made a broad head-turn that encompassed the whole range of directions of the cell’s whole-trial-averaged tuning. If the broader tuning curves were caused entirely by attractor drift, this instantaneous estimate will be narrower than the trial-averaged tuning curve, and not different between lesion and control animals. Indeed, in both Sham and Lesion HD cells, the mean width of the instantaneous tuning curve was smaller than the tuning width averaged across the whole trial (Sham: 44.3 ± 33.0°, Lesion: 44.9 ± 34.6°, Figure 8C). No significant difference in instantaneous tuning curve width was observed between groups (z = 0.634, p = 0.526), indicating that the wider tuning widths in Lesion animals occur primarily due to attractor drift across the trial.

Having confirmed that tuning curves in Lesion animals drifted more across a trial, we then looked to see whether multiple simultaneously recorded cells drifted together. We looked at co-recorded cell pairs in short time windows of 60 seconds. Each window was step-wise advanced across the recording in 12-s increments, the mean spiking direction calculated for each cell in that window and then FDs pairwise compared by computing a Rayleigh vector of the angular differences. Relative angles between co-recorded HD cells remained in close accordance throughout trials in both Lesion and Sham animals (mean HD cell pair R value per session: Lesion: 0.81 ± 0.17, Sham: 0.74 ± 0.12, Figure 8D). Lesion HD cells were significantly more coherent than Sham HD cells at the session level (t(136) = 2.60, p = 0.01), although no significant difference was observed at a per-animal level (t(8) = −1.16, p = 0.28).

#### Anchoring of HD cells to visual polarising cues was impaired in animals with dLGN lesions

Having looked at within-trial stability, we then turned to the question of between-trial stability and the extent to which firing directions were controlled by the visual landmarks. An example recording of an HD cell recorded from a Sham animal is shown in Figure 9A. Sham HD cells showed good visual landmark anchoring to all tested cue cards, as evidenced by the normalised FD deviations from expected which were clustered around zero. This was shown by a significant Rayleigh vector (*R* = 0.77, p < 10^-53^, n = 169 trial shifts (Figure 9B) which on testing showed to be towards the direction zero (V0° = 130.4, p < 10^-51^). The mean absolute deviation from the expected shift (0° if the cells were anchored to the cue cards) was 26.7° ± 39.5°.

**Figure 9:**
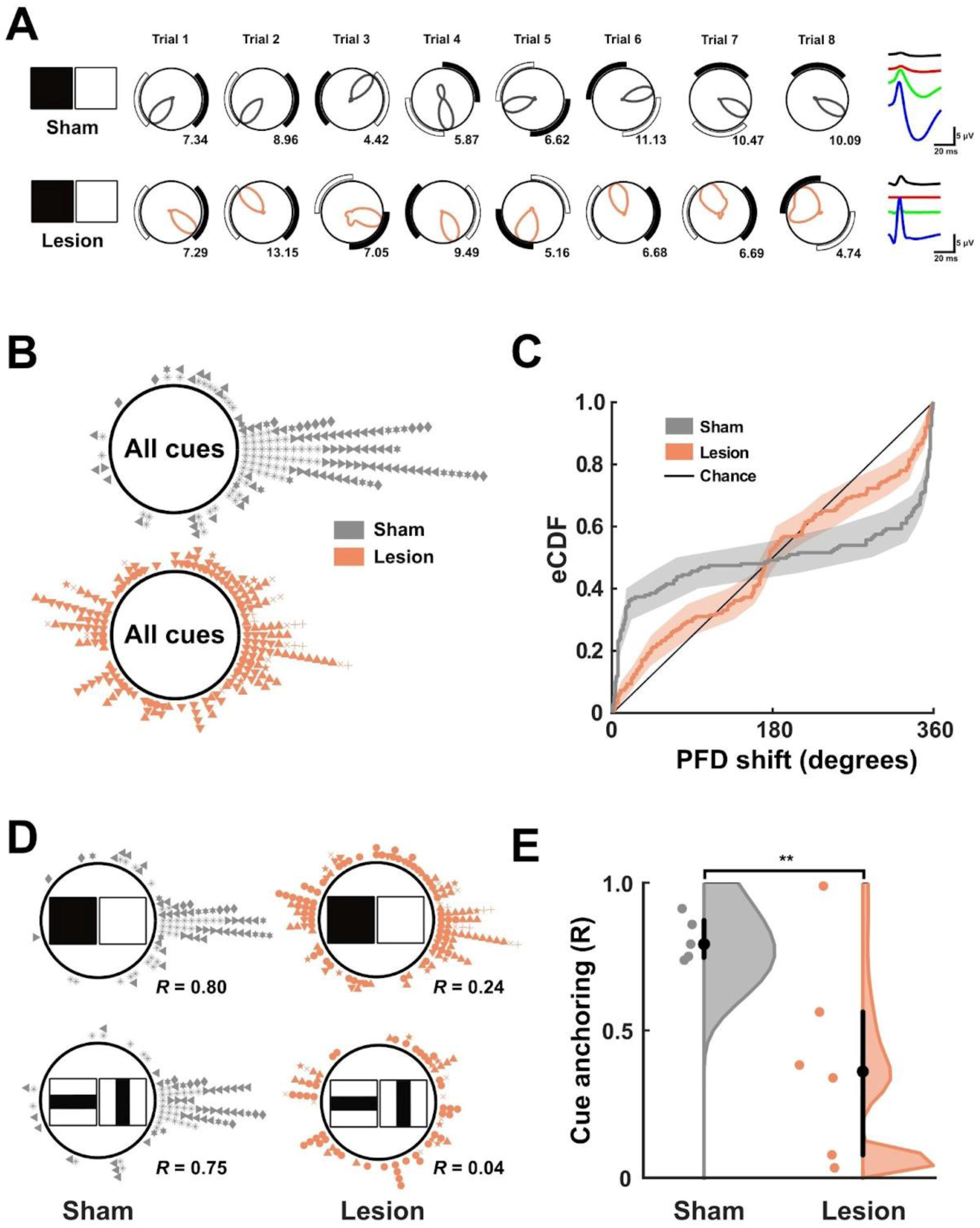
Landmark anchoring of HD cells in Sham and Lesion animals. A: Example sessions showing a cell from a Sham and Lesion animal (see waveforms at far right) in response to shifts of the black and white cue cards. Note that the tuning curve was slightly broader in the cell from the lesion animal, and that it did not reliably follow the cue. B: Summary of all FD shift errors in Lesion (top, orange) and Sham (bottom, grey) animals across all trials in black-white and vertical-horizontal cue conditions. Accurate landmark anchoring is represented by a 0° shift error (positioned right on the circle). C: Cumulative distribution of all PDF shift errors in Lesion (orange) and Sham (grey) animals across all trials in black-white and vertical-horizontal cue conditions. D: Summary of FD shift errors in Lesion (orange) and Sham (grey) animals across all trials separated by cue condition (black-white, top and vertical-horizontal, bottom). E: Landmark anchoring success across all Lesion (orange) and Sham (grey) animals.

Across Sham sessions, we saw no evidence of any difference in anchoring precision to black-white versus vertical-horizontal cue cards (RBW = 0.80, p < 10-29; RVH = 0.75, p < 10^-24^; Kuiper test: k = 1235, p > 0.5), with similar absolute deviations from expected shifts (BW: 24.3° ± 34.7°; VH: 29.1° ± 44.0°). Overall, these results suggest that both black-white and vertical-horizontal cue conditions provide salient, precise visual landmarks that can be used by the HD system for anchoring.

Contrastingly, landmark anchoring in Lesion animals was significantly impaired relative to Sham. Over all tested cue cards, there was only weak evidence of non-uniformity in normalised FD shifts (*R* = 0.18, p < 0.005, n = 194 trial shifts). This was, however, towards the direction 0° (V0° = 34.2, p < 0. 001), indicating that some weak landmark anchoring was present. However, there was more variance in the shifts, with the absolute deviation from expected shift being 76.5° ± 62.7°.

In Lesion animals there was also a difference in precision of landmark anchoring to the different cue-card pairs. While some residual anchoring was observed to the high-contrast black-white pair (*R* = 0.240, p < 0.001; absolute deviation = 71.2° ± 63.4°), there was no evidence of successful anchoring to the low-contrast vertical-horizontal pair (*R* = 0.09, p = 0.58; absolute deviation = 86.0° ± 59.9°). The distributions of FD shifts to these sets of cue cards (Figure 9C) were significantly different (Kuiper test: k = 2479, p < 0.05). Lesion landmark anchoring was significantly impaired relative to Sham in all non-ambiguous cues (k = 14598, p < 0.001), black-white cues (k = 4524, p < 0.001), and vertical-horizontal cues (k = 3584, p < 0.001).

A summary statistic was derived corresponding to the anchoring success in each animal (see Methods); average anchoring in Lesion animals to non-ambiguous cues was significantly impaired relative to Sham (mean R vector, Lesion animals: 0.398 ± 0.351, n = 6; Sham animals: 0.811 ± 0.074, n = 5; t = −2.56, p = 0.03; Figure 9E). Importantly, these results cannot be explained due to the averaging of FD shifts of multiple HD cells in a drifting attractor: a per-animal anchoring statistic using only the single “best anchored” HD cell in each session was also significantly reduced in Lesion animals (Lesion: 0.456 ± 0.284, n = 6; Sham: 0.820 ± 0.046, n = 5; t = 2.81, p = 0.0205), indicating that the impairment was present at the individual cell as well as at the population level.

#### Lesion and Sham HD cells did not use non-visual information to anchor to landmarks

Disruptions of the visual system in Lesion animals may increase reliance on non-visual orienting cues (for example to the smell of the cue cards), which could explain the presence of weak residual landmark anchoring. The two-fold visual symmetry of the environment allowed us to decouple detection of the presence of cue cards (for instance, through whiskering against its edges along a wall) from discrimination of their visual content. An animal that can detect, but not visually discriminate, the cue cards will show a bipolar distribution of FD shifts across sessions. The way to reveal this is to show that the Rayleigh vector of the FD shifts is much greater if the angles are doubled, causing the values at 180 degrees to map to 0/360 while those at 0/360 still map to 0/360.

We saw such a bipolar distribution in Sham animals tested in the identical-cue condition. This was shown by a low resultant vector for the raw data (*R* = 0.075, p = 0.70, n = 63 shifts) coupled with a much larger R vector for the angle-doubled data (Sham: angle-doubled *R* = 0.816, p < 10^-22^). This indicates that Sham HD cells could use the cue cards to anchor, but could not distinguish them and thus were not using non-visual cues. A similar distribution was seen in Lesion animals for black-black FD shifts (*R* = 0.063, p = 0.71, n = 84 shifts), which also increased following angle-doubling (Lesion: angle-doubled *R* = 0.620, p < 10^-15^). The change in *R* was greater in Sham animals than Lesion animals (Sham: 0.74; Lesion: 0.56), with angle-doubled Lesion FD shifts remaining significantly less precise than angle-doubled Sham shifts (Kuiper test: k = 1932, p = 0.002).

We next looked at the patterned cues (Fig. 9D), starting with the high-salience black-white cues. Here, in Sham animals we saw (as expected) the reverse pattern in which *R* values were high initially (*R* = 0.800), and declined slightly on angle-doubling (*R* = 0.609, p < 10^-15^) which was significantly lower (Kuiper test, k = 1955, p < 0.05). This indicates that the distribution of FD shifts was predominantly unimodal. By contrast, across Lesion sessions, initial *R* values were low (*R* = 0.240) and increased on angle-doubling (angle-doubled R = 0.422, p < 10^-10^; Kuiper test, k = 3668, p < 0.05) suggesting that the cues were detectable but not discriminable. A similar pattern of results was seen in vertical-horizontal sessions: the resultant vector decreased in magnitude following angle-doubling in Sham animals (from *R* = 0.745 to *R* = 0.669) but increased in Lesion animals (from *R* = 0.037 to *R* = 0.273) although it remained somewhat low.

Together, these results show that HD cells in Sham animals could discriminate the visually different cues but not the visually identical ones, ruling out non-visual use of the cues (olfaction etc.) and confirming the use of pattern vision. Lesion animals by contrast showed some evidence of cue use but failed to use either luminance (black vs. white cards) or pattern, based on their failure to distinguish these cues, although the black-white cues did exert slightly better control than the more subtly different patterned cues. The residual cue use suggests that residual anchoring may possibly be contributed to by symmetric non-visual information such as tactile content.

## Discussion

We investigated whether the retino-geniculo-cortical visual pathway is necessary for the responsiveness of rodent HD cells to visual landmarks, or whether this responsiveness could be supported by the retino-tectal pathway alone. In dLGN-lesioned rats we recorded HD cells from the PoS, which is a cortical region located within the visual system, observing apparently normal numbers of HD cells here. These had broader whole-trial tuning curves, but we found that this was due to the drift of tuning curves across a trial. When we corrected for this we found that the underlying tuning curves were as precise as in control rats and showed normal coherence between pairs of neurons. Despite this local tuning precision, visual landmark cues failed to exert control over the neurons in the lesioned animals, even though these cues were large and high-contrast. These results indicate that the subcortical retino-tectal pathway, despite being by far the largest visual pathway on rats, is insufficient to provide visual anchoring of the HD system to the external world even for coarse high-contrast stimuli. This is surprising because this pathway is phylogenetically much older than the neocortex, and likewise, visual determination of heading direction is an ancient and fundamental competence, which might have been expected to precede cortical visual landmark processing. Indeed, many species lacking a cortex can use visual scene for orientation (Schwarz *et al*., 2024). Below, we explore the nature and implications of these findings.

### dLGN lesions mildly increased exploratory behaviour

We first examined whether behavioural measures such as speed of movement, speed of turning, turning bias or thigmotaxis (wall-hugging) might be altered in lesioned animals, since vision is important for movement-planning, and movement parameters can affect HD cell tuning curves. For example, thigmotaxis might result in restricted directional sampling, degrading tuning curves for the under-sampled directions. However we did not see notable changes in these parameters. Lesioned animals did show a mild increase in exploratory behaviour, revealed by a slight increase in total distance travelled and total cumulative angular head turning, but these particular parameters should not have affected tuning curve width or stability. Interestingly, coverage of the arena was the same in both lesioned and sham groups, with both groups spending almost 50% of their time in the central 50% of the arena, suggesting that the lesioned animals were not anxious about venturing away from the tactile self-localisation afforded by the walls. We assume this was not due to residual vision, because of the lesion effect on landmark responsiveness we saw in the neurons. It might instead be because olfactory cues across the floor were able to provide sufficient information for self-localisation. A previous experiment recording place cells in blind rats found that place field precision remained high if olfactory cues on the floor were preserved but not if they were disrupted (Paz-Villagràn *et al*., 2002). Overall, our behavioural observations did not find any lesion effects that might have altered HD cell firing characteristics.

### Intact basic directional firing and firing coherence in dLGN-lesioned animals

Our next question was whether we would find HD cells at all, in animals deprived of geniculo-cortical vision. We found them in seemingly normal numbers, and with normal firing rates. This is as expected, because HD cells are found even in newborn rat pups prior to eye opening (Bjerknes *et al*., 2015; Tan *et al*., 2015) and also in mice that have lost vision due to retinal degeneration (Asumbisa *et al*., 2022). This observation adds weight to the notion that HD tuning curves are generated independently of visual input, via a pathway that starts in the vestibular nuclei and progresses through a series of brainstem nuclei to eventually reach the anterior thalamus (Cullen & Taube, 2017).

The fact that HD tuning curves are well-formed and stable even in animals deprived of vision indicates that other senses are contributing significantly. One of these could be olfaction, which as mentioned above is able to support place cell firing in blind animals. With HD cells, it has been consistently found that the cells under-rotate their tuning curves when a visual landmark is rotated (see (Knight *et al*., 2014) for data and review), possibly due to a counteracting influence of unrotated olfactory cues (although other background cues could also contribute). More recently, Asumbisa et al. (2022) showed that olfactory information is critical for HD cell stability as olfactory ablation resulted in complete loss of HD cell tuning in the anterior thalamus of blind mice. When olfactory information is present, however, then the signal retains a reasonable degree of stability despite the increased variability described above.

We also found that despite an increased rate of drift (discussed below) the coherence between simultaneously recorded cells remained intact. Asumbisa et al. also saw this in their retinal degeneration study although these HD cells were recorded in the thalamus (not cortex) of mice (not rats). If HD cells act independently then the noisy firing reflected in the increased drift rate should result in reduced coherence (more variable angular distances between pairs of tuning curves). That this was not observed suggests that the cells are *not* acting independently, and is consistent with attractor models of HD firing (Angelaki & Laurens, 2020) in which firing directions are linked by virtue of the cells’ interconnections. Our findings thus imply, as Asumbisa et al. also concluded, that attractor dynamics are not dependent on cortical visual input. This is unsurprising, since the attractor has long been thought to be located subcortically, perhaps between the dorsal tegmental and lateral mammillary nuclei in the brainstem (Taube, 2007). Firing rates were also not different in lesioned animals despite the absence of one of the usual sources of incoming sensory information. This is also typical for HD cells (for example they do not reduce their firing in the dark) and is presumably because these external sensory sources do not drive the system so much as orient it, by biasing activity in the attractor network towards a given part of it.

### Broader HD cell tuning curves in dLGN-lesioned animals that drifted across a trial

One difference we did find in lesioned animals is that tuning curves were 20% broader when the whole-trial average was analysed in the usual way. In order to determine whether this could be due to a broader instantaneous attractor bump (region of activation) or to a normal tuning curve that was more unstable, we measured the instantaneous width of activation as closely as possible by examining only those segments of firing where a given head turn spanned the entire region of the (broadened) tuning curve. We found that the width of the firing region now did not differ between lesioned animals and controls. Thus, the broadened tuning curve is due to a narrow tuning curve that wobbles. This is consistent with the anchoring of the tuning curve having been deprived of the more precise orientation provided by visual cue, relying instead on a combination of self-motion cues and olfaction. The trial-averaged drift rate (linear drift rate of 7°/min, mean shift of 30° in 4 minutes) in lesioned animals was found to be similar in magnitude to that reported previously for blindfolded rats: 2.9°/min (Goodridge *et al*., 1998).

### Anchoring of HD cells to visual polarising cues was impaired in animals with dLGN lesions

A notable difference that we found between dLGN-lesioned animals and controls is that in the lesioned animals, PoS HD cells failed to respond to reorientation of polarising visual cues, distributing their responses more or less randomly around the compass circle rather than clustering near the position signalled by the cues. This was the case not just for the cue-pairs that differed in their details, but also the back-white cue pair for which the cues were large, high-contrast and very distinctive. By contrast, the sham animals showed good clustering even for the harder-to-distinguish cues. It thus seems that polarising visual information did not reach the PoS HD cells, indicating that the intact collicular pathway provided insufficient visual information about the landmarks.

On the one hand this seems surprising: in rodents, the retino-tectal pathway is much larger than the geniculo-cortical one, with 85-90% of retinal ganglion cells projecting into superior colliculus (Ellis *et al*., 2016). It is phylogenetically older (Knudsen, 2020), and so plausibly may have provided a primordial HD signal in the ancestral forerunner to mammals. However, there is no evidence that superior colliculus performs parametric analysis of visual features (Knudsen, 2020). Rather, neurons in the superficial superior colliculus of rodents encode various behaviourally-relevant stimulus characteristics, including looming, fast- and slow-moving stimuli (Gale & Murphy, 2014), and display reasonable spatial acuity, albeit less fine-tuned than neurons in V1 (Li et al., 2015).

Consistent with this, there are no reports, to our knowledge, of direct connectivity between superior colliculus and any of the HD areas. A possible indirect route exists via the postrhinal cortex, in which activity has been shown to critically depend exclusively upon inputs from superior colliculus, but not V1 (Beltramo & Scanziani, 2019). However, lesions of the postrhinal cortex do not disrupt landmark anchoring in HD cells (Peck & Taube, 2017). The geniculo-striate pathway, by contrast, is more plausibly involved in landmark control of HD cells. Both V1 and extrastriate areas directly project into PoS and retrosplenial cortex (van Groen & Wyss, 1990; Wang *et al*., 2012), and lesion studies of landmark control suggest that landmark information enters the HD circuit at the level of these cortical areas (Clark *et al*., 2010)(Yoder *et al*., 2015). The retrosplenial cortex in particular contains landmark-sensitive multi-directional cells (Jacob *et al*., 2017; Zhang *et al*., 2022).

In sum, our present data add to this body of literature by providing the first direct evidence that it is the geniculo-striate pathway that provides landmark information to HD cells, and therefore likely to downstream spatial circuits, strongly suggesting landmark control of directional orientation is predominantly supported by cortical vision in rodents.

## Conclusion

This study has shown that the dLGN is required for landmark control of cortical (and presumably other) head direction cells, but is not required for the generation of directionally precise tuning curves per se. This is the case even though the retino-tectal pathway is by far the largest visual pathway in rodents, and evolved long before the geniculo-cortical pathway. These findings raise questions about the evolution of directional landmark processing in vertebrates, and the extent to which animals lacking a cortex can process the structure of visual landmarks for use in directional orientation. It may be that they cannot, or alternatively that in mammals, the geniculo-cortical pathway has taken over a function that earlier in evolution was supported by the optic tectum

## Acknowledgements

**Author contributions:** James Street: Conceptualization, Methodology, Software, Investigation, Formal analysis, Data curation, Visualization, Writing – Original Draft, Review & Editing; Kate Jeffery: Conceptualization, Methodology, Data curation, Supervision, Resources, Funding acquisition, Writing – Review & Editing. **Competing interests**: The authors declare the following competing interests: K. J. J is a non-shareholding director of Axona Ltd.

## Data and materials availability

Data and MATLAB code used for analysis and figure generation are available at the UCL Research Data Repository (doi: 10.5522/04/25476172), and curated MATLAB code will be made available at the author’s GitHub (https://github.com/bejamz).

